# LncPNdeep: A long non-coding RNA classifier based on Large Language Model with peptide and nucleotide embedding

**DOI:** 10.1101/2023.11.29.569323

**Authors:** Zongrui Dai, Feiyang Deng

## Abstract

Long non-coding RNA plays an important role in various gene transcription and peptide interactions. Classifying lncRNAs from coding RNA is a crucial step in bioinformatics analysis which seriously affects the post-analysis for transcriptome annotation. Although several machine learning-based methods were developed to classify lncRNAs, these methods were mainly focused on nucleotide features without considering the information from the peptide sequence. To integrate both nucleotide and peptide information in lncRNA classification, one efficient deep learning is desired. In this study, we developed one concatenated deep neural network named LncPNdeep to combine this information. LncPNdeep incorporates both peptide and nucleotide embedding from masked language modeling (MLM), being able to discover complex associations between sequence information and lncRNA classification. LncPNdeep achieves state-of-the-art performance in the human transcript database compared with other existing methods (Accuracy=97.1%). It also exhibits superior generalization ability in cross-species comparison, maintaining consistent accuracy and F1 scores compared to other methods. The combination of nucleotide and peptide information makes LncPNdeep able to facilitate the identification of novel lncRNA and gain high accuracy for classification. Our code is available at https://github.com/yatoka233/LncPNdeep

## 1. Introduction

Long non-coding RNA (lncRNAs) is a heterogeneous class of non-coding RNA with nucleotide lengths larger than 200bp [1]. LncRNA can regulate the gene expression and developmental process which involves X chromosome inactivation, genomic imprinting, and Hox gene regulation [2–5]. In human diseases, lncRNAs play a crucial role in breast cancer progression, hepatocellular carcinoma, and Alzheimer’s disease [6–8]. Identifying and understanding the functions of lncRNAs is vital in genetic studies.

Previous lncRNA classification methods are mainly based on deep learning and machine learning strategies. These methods utilized the features extracted from nucleotide sequences to make predictions. For instance, CPC2 applied a support vector machine (SVM) with four intrinsic features extracted by the random forest model [9]. PLEK designed one improved k-mer feature to build the SVM model for prediction [10]. The Coding-Potential Assessment Tool (CPAT) was another alignment-free model that utilized logistic regression with the Fickett Score, Hexamer Score, ORF features to identify novel lncRNAs [11]. CNCI made use of the frequency of ANT in the coding region to design features. Using a dynamic programming algorithm to find the Most-Like Coding Domain Sequence (MLCDS) and incorporate five features from it for SVM model training [12]. FEELnc developed a pipeline including filtering and classifier together for prediction. It designed multi k-mer frequency and ORF coverage proportion as input of their random forest model [13]. Also, other machine learning such as stacked ensemble models and neural networks were also applied in lncRNA identification [14,15]. The above methods were machine learning-based methods with nucleotide features as their input. While several methods were mainly based on deep learning structure for prediction. For instance, DeepPlnc was one bi-model CNN deep learning model which can utilize nucleotide and structural information parallelly [16]. PlncRNA-HDeep used Long short-term memory (LSTM) and Convolution neural network (CNN) to extract the abstract features from the nucleotide sequence, avoiding complex feature selection [17]. lncRNA_Mdeep was one multimodal deep learning structure that concatenate OFH features (ORF feature, Fickett score, and Hexamer score), k-mer features, and one-hot coding of sequence together [18]. lncRNA_MFDL is one deep stacked neural network (DSN) that fused the ORF, k-mer feature, and coding domain sequence features together [19]. However, all these models only considered the nucleotide features extracted from sequences without considering the information from their peptide sequences.

Some studies found that certain lncRNAs that existed in human cells may have the ability to translate themselves. While most of the translated peptides generated from lncRNAs were highly unstable and could not function normally [20]. Also, few lncRNAs can translate into short peptides that do have proper biological functions [21–23]. These lncRNAs may consist of short open reading frames (sORFs) and have weak coding ability [24]. This information indicates that the possible peptide from lncRNA’s ORF may have certain functions or structures which is different from normal protein. To capture the peptide and nucleotide together, we utilized several large language models to extract the information. It’s an important aspect to incorporate nucleotide information robustly without extracting complex features or calculating scores. This notion is inspired by the intriguing parallelism between RNA sequences and natural languages due to their shared sequential structures [25,26]. This potential cross-disciplinary endeavor allows us to transpose sequence characterization methodologies from the field of NLP into lncRNA classification. In the domain of Natural Language Processing, large language models have risen to prominence, consistently delivering state-of-the-art results in a wide range of tasks, including sentence classification and question answering [27,28]. Research has shown that the incorporation of self-supervised and semi-supervised tasks during the pre-training phase can enhance the acquisition of generalized and robust features within the latent space of raw sequences [29–31].

To capture the peptide and nucleotide information together, we utilized large language models to extract the peptide and nucleotide information. The peptide embedding is extracted by the ProtTrans model which is a large language model trained by self-supervised learning [32]. Bibird and Longformer models were used to pre-train RNA sequences and obtain the nucleotide embeddings. Based on the embeddings, we proposed one concatenated deep learning method, LncPNdeep, to classify lncRNAs based on the peptide and nucleotide embeddings. LncPNdeep mainly contains three parts: (1) Convolution 1D layer with a residual connection that takes peptide embedding from ProteinTrans as input; (2) Convolution 1D layer with a residual connection that takes nucleotide embedding from Longformer and Bigbird as input; (3) Bidirectional LSTM layers that learning the concatenated information.

## 2. Materials and Methods

### 2.1. Dataset

The coding RNA and lncRNA for humans were downloaded from the GENCODE Release 43 (GRCh38.p13) [33]. The original data contained 54,291 lncRNA and 110,224 coding RNA. The training dataset contains 48,876 lncRNAs and 99,187 Coding RNAs. The test dataset contains 5,415 lncRNAs and 11,037 Coding RNAs (Table 1). In the training dataset, we applied bootstrapping to resample the lncRNAs 99,187 times. Making training samples balanced. Cross-species datasets contained 6 vertebrates and 6 fungi. Vertebrates’ lncRNAs and coding RNAs were downloaded from the ensemble database [34]. The vertebrate data consists of Pandas (*Ailuropoda melanoleuca*), Cows (*Bos taurus*), Zebrafish (*Danio rerio*), Chicken (*Gallus gallus*), Gibbons (*Nomascus leucogenys*), and Pig (*Sus scrofa*). The fungi datasets were downloaded from the ensemble fungi database. It consists of *Aspergillus nidulans, Magnaporthe grisea, Puccinia graminis, Saccharomyces cerevisiae, Schizosaccharomyces pombe, Zymoseptoria tritici*.

**Table 1.** The human RNA dataset separation for training and testing.

|  | LncRNAS | Coding RNAs |
| --- | --- | --- |
| Train dataset | 48,876 | 99,187 |
| Test dataset | 5,415 | 11,037 |
| Total | 54,291 | 110,224 |

### 2.2. Peptide embedding

We applied the ProtTrans model, a large language model to extract the peptide information from coding RNAs and lncRNAs. Since some lncRNAs didn’t contain ORFs, we designed three peptide embeddings named Fake peptide embedding, Max peptide embedding, and Average peptide embedding. In the first step, the algorithm will search the start codon (ATG) and stop codon (TAG, TGA, TAA). If no ORFs are detected, the first three bases will be taken as the first codon and start the translation. This peptide will be passed to the ProtTrans model and obtain its embedding – Fake peptide embedding. If at least one ORF is detected, the ORF with maximum length will be passed to the ProtTrans model and obtain the Max peptide embedding. Average peptide embedding was computed by the average value of multiple peptide embedding. If the nucleotide sequence doesn’t have ORF, the Max peptide and Average peptide embedding will be replaced by the Fake peptide embedding.

### 2.3. Nucleotide embedding

Numerous lncRNAs exhibit extensive lengths which surpasses the capacity of conventional transformer-based models for processing. Many works have investigated approaches to long sequences, and in this study, we harness the capabilities of the BigBird and Long-former models, renowned for their efficacy in handling lengthy sequences [35–39]. We consider each base of the sequence as a token so that our approach can also entail refinements to the existing Masked Language Model (MLM) pre-training framework by adding a [*CLS*] token at the beginning of all sequences and replacing 15% of the tokens with [*MASK*] tokens. Taking inspiration from the genetic manipulation techniques employed in the BigBird model, we address the challenge posed by exceedingly protracted RNA sequences. For each RNA sequence, we adopt a strategy of sampling k subsequences to represent it, whose lengths are within the defined range [*l, u*]. The length of the subsequences is thus limited to the maximum input length of the model. During the pre-training phase, our focus lies solely on training the sampled subsequences with the MLM loss, i.e., identifying the original tokens on the masked positions. Subsequently, during the prediction phase, the output embedding of the [*CLS*] tokens is used as the subsequence representation. In order to predict the coding status, an aggregation (here we use average) of output representations across all the identified subsequences for each RNA sequence is generated. In our setting, three types of embeddings are generated for each RNA sequence, i.e., Longformer256 embedding, Bigbird256 embedding, and Bigbird768 embedding.

### 2.4. Concatenated deep learning

After obtaining the peptide embeddings (Fake peptide embedding, Max peptide embedding, and Average peptide embedding) and nucleotide embeddings (Longformer256, Bigbird768, Bigbird256), each embedding will be passed through one customed CNN model which contains four Convolution 1D layers with residual connections. The outputs are then merged to form a comprehensive feature map with 4352 dimensions. To prevent overfittings, the final feature map would be passed through a dropout layer (Dropout rate=0.8). Two BiLSTM layers extract the information from the filtered feature map and obtain the classification result through one DNN layer. The Categorical Cross entropy is taken as the loss function of LncPNdeep.

## 3. Results

To evaluate the prediction performance of lncPNdeep and other existed tools, we applied our model under human transcriptome and cross-species datasets. Existed methods including CNCI, CPC2, CPAT, PLEK, LncAdeep, and LncRNA_Mdeep were compared with lncPNdeep in human RNA datasets. To benchmark these tools, the best way is to re-train these models on the same training dataset and test their performance on the unique test dataset. For the models without retrain options such as CNCI, CPC2, LncAdeep, and LncRNA_Mdeep, we used their pre-trained model directly. Among PLEK and CPAT models, the retraining was very time-consuming due to the large sample size of our training dataset. So, we used the pre-trained version of these methods for human transcriptome comparisons.

### 3.1. Performance of human lncRNA identification

To validate whether our model can outperform the traditional machine learning algorithm, we compared LncPNdeep with Support Vector Machine (SVM), Logistic Regression (LR), and Random Forest (RF). Each baseline model used the same training and testing datasets in the human transcriptome. Since each sample had six embeddings: peptide embeddings (Max, Average, and Fake embeddings) and nucleotide embeddings (Long-former256, Bigbird256, and Bigbird768), each baseline model used one embedding for training. Compared to the performance of each baseline model in the test dataset, the LncPNdeep model had superior accuracy, precision, and recall value on the test dataset. Among the baseline models, SVM with Bigbird256 embeddings as a training dataset had the highest accuracy value compared to other models (Accuracy = 0.93). In each baseline machine learning algorithm, the model with Bigbird256 embeddings achieved the best performance compared to others (Table 2). We also compared the performance of LncPNdeep with the other 6 existing methods for lncRNA classification on the human RNA dataset. In the result below, LncPNdeep achieved 97.1% accuracy, 96.7% specificity, and 98.0% sensitivity (Table 3, etc.). Its sensitivity ability outperforms all the existing methods which were 3.6%, 7.0%, 27.3%, 17.5%, 64.8%, and 28.4% higher than LncAdeep, LncRNA_Mdeep, CPC2, CNCI, CPAT, and PLEK. LncAdeep had the closest performance with LncPNdeep which achieved 95.1% accuracy, 96.2% specificity, and 94.4% sensitivity. CPAT had the worst accuracy in the human dataset (accuracy = 53.9%). This result may be due to its poor sensitivity in lncRNA classification (sensitivity = 33.2%).

**Table 2.** The performance comparisons in the human test dataset with baseline models.

| Model | Feature Embedding | Accuracy | Precision | Recall |
| --- | --- | --- | --- | --- |
| **LncPNdeep** | All | **0.95** | **0.90** | **0.95** |
| **SVM** | Max Peptide | 0.80 | 0.67 | 0.75 |
|  | Aver Peptide | 0.75 | 0.62 | 0.63 |
|  | Fake Peptide | 0.82 | 0.70 | 0.77 |
|  | Bigbird256 | 0.93 | 0.86 | 0.94 |
|  | Bigbird768 | 0.87 | 0.76 | 0.89 |
|  | Longformer256 | 0.87 | 0.79 | 0.81 |
| **RF** | Max Peptide | 0.83 | 0.75 | 0.73 |
|  | Aver Peptide | 0.84 | 0.76 | 0.74 |
|  | Fake Peptide | 0.84 | 0.76 | 0.74 |
|  | Bigbird256 | 0.92 | 0.86 | 0.92 |
|  | Bigbird768 | 0.88 | 0.80 | 0.86 |
|  | Longformer256 | 0.86 | 0.81 | 0.73 |
| **LR** | Max Peptide | 0.81 | 0.70 | 0.73 |
|  | Aver Peptide | 0.79 | 0.67 | 0.73 |
|  | Fake Peptide | 0.83 | 0.73 | 0.75 |
|  | Bigbird256 | 0.92 | 0.86 | 0.92 |
|  | Bigbird768 | 0.88 | 0.79 | 0.85 |
|  | Longformer256 | 0.85 | 0.78 | 0.76 |

**Table 3.** The performance comparisons in the human dataset with existing methods.

|  | Accuracy | Specificity | Sensitivity |
| --- | --- | --- | --- |
| LncPNdeep | **0.971** | **0.967** | **0.980** |
| LncAdeep | 0.951 | 0.962 | 0.944 |
| LncRNA_Mdeep | 0.922 | 0.944 | 0.910 |
| CPC2 | 0.781 | 0.921 | 0.707 |
| CNCI | 0.855 | 0.951 | 0.805 |
| CPAT | 0.539 | 0.965 | 0.332 |
| PLEK | 0.771 | 0.914 | 0.696 |

### 3.2. Performance of cross-species lncRNA identification

To testify whether LncPNdeep had generalization ability to cross-species datasets, we applied the LncPNdeep method in 12 species RNA databases. With only human RNA as the training dataset, LncPNdeep demonstrated remarkable accuracy and F1 in most of the species except the Cow, *S. pombe*, and *Z. tritici*. Specifically, LncPNdeep achieved an accuracy of 95.5% and an F1 score of 97.6% in pandas, 93.5% accuracy and 97.7% F1 score in cows, and 90.6% accuracy with a 96.3% F1 score in zebrafish (Table 4). In the existing methods, CPAT had the best accuracy on Cow (accuracy = 95.0%) and *Z. tritici* (accuracy = 95.7%) while CPC2 obtained the best accuracy in the *S. pombe* (accuracy = 94.1%). LncPNdeep consistently maintained high F1 scores across all species which never dropped below 90%, reflecting its robustness and reliability in cross-species RNA analysis.

**Table 4.**
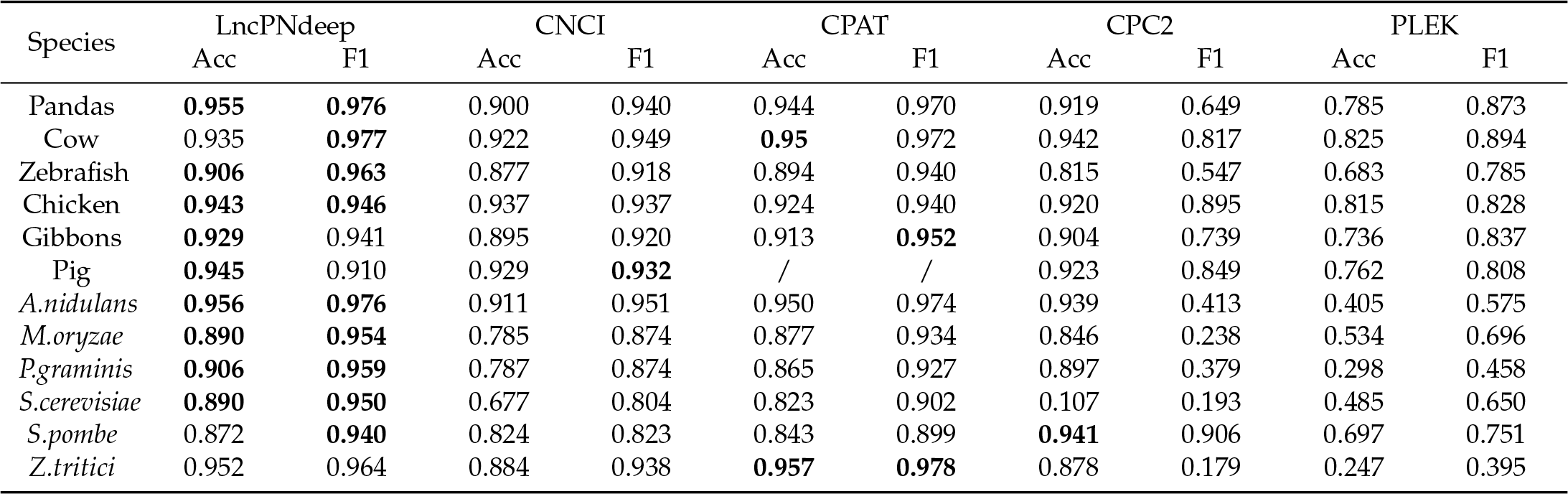
The performance of LncPNdeep and other four methods on 12 cross-species datasets.

| Species | LncPNdeep |  | CNCI |  | CPAT |  | CPC2 |  | PLEK |  |
| --- | --- | --- | --- | --- | --- | --- | --- | --- | --- | --- |
|  | Acc | F1 | Acc | F1 | Acc | F1 | Acc | F1 | Acc | F1 |
| Pandas | **0.955** | **0.976** | 0.900 | 0.940 | 0.944 | 0.970 | 0.919 | 0.649 | 0.785 | 0.873 |
| Cow | 0.935 | **0.977** | 0.922 | 0.949 | **0.95** | 0.972 | 0.942 | 0.817 | 0.825 | 0.894 |
| Zebrafish | **0.906** | **0.963** | 0.877 | 0.918 | 0.894 | 0.940 | 0.815 | 0.547 | 0.683 | 0.785 |
| Chicken | **0.943** | **0.946** | 0.937 | 0.937 | 0.924 | 0.940 | 0.920 | 0.895 | 0.815 | 0.828 |
| Gibbons | **0.929** | 0.941 | 0.895 | 0.920 | 0.913 | **0.952** | 0.904 | 0.739 | 0.736 | 0.837 |
| Pig | **0.945** | 0.910 | 0.929 | **0.932** | / | / | 0.923 | 0.849 | 0.762 | 0.808 |
| *A.nidulans* | **0.956** | **0.976** | 0.911 | 0.951 | 0.950 | 0.974 | 0.939 | 0.413 | 0.405 | 0.575 |
| *M.oryzae* | **0.890** | **0.954** | 0.785 | 0.874 | 0.877 | 0.934 | 0.846 | 0.238 | 0.534 | 0.696 |
| *P.graminis* | **0.906** | **0.959** | 0.787 | 0.874 | 0.865 | 0.927 | 0.897 | 0.379 | 0.298 | 0.458 |
| *S.cerevisiae* | **0.890** | **0.950** | 0.677 | 0.804 | 0.823 | 0.902 | 0.107 | 0.193 | 0.485 | 0.650 |
| *S.pombe* | 0.872 | **0.940** | 0.824 | 0.823 | 0.843 | 0.899 | **0.941** | 0.906 | 0.697 | 0.751 |
| *Z.tritici* | 0.952 | 0.964 | 0.884 | 0.938 | **0.957** | **0.978** | 0.878 | 0.179 | 0.247 | 0.395 |

### 3.3. Importance of peptide embedding towards prediction

We conducted a permutation experiment to determine the contribution of each peptide embedding to the predictive ability of the LncPNdeep model. For each permutation, we randomized one set of peptide embeddings in a human test dataset and evaluated the model’s performance. This process was repeated 300 times to ensure reliability. The resulting violin plots illustrated the distribution of the model’s performance metrics across these permutations (Figure 1). It is evident that the permutation of peptide embeddings leads to a varied distribution of metrics, confirming the significance of peptide embeddings in the model. Among the three peptide embeddings, Average peptide embeddings contributed most to the predictive ability which has the lowest mean of accuracy and specificity after randomization.

**Figure 1.**
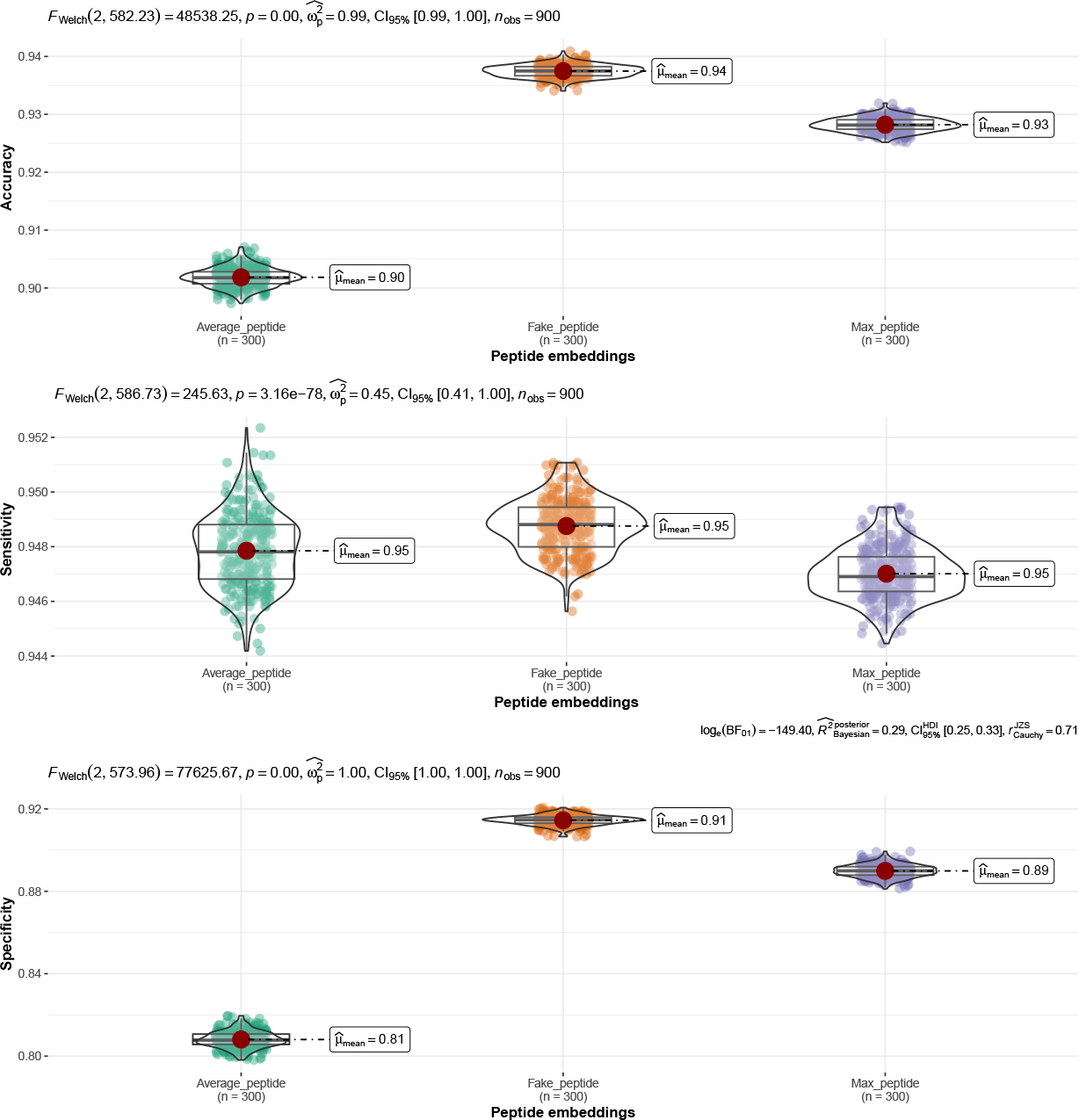
Permutation experiments of lncPNdeep model.

### 3.4. t-SNE clusters of each embedding

To determine the performance of each individual and combined embeddings, we utilized t-SNE clustering on each embedding to visualize their separation. All the embeddings showed a noticeable overlapping between coding RNA and lncRNA. In Bigbird768 and Longformer256 embeddings, the clusters were mixed up with each other which means a solely nucleotide embedding was not sufficient for classification. In the peptide and Bigbird256 embeddings, they had slightly better separation of lncRNA and coding RNA, but the overlapping remained substantial (Figure 2). These results indicated that only based on a single nucleotide or peptide embedding was not ideal for lncRNA classification. Therefore, employing combination or ensemble techniques to amalgamate these embeddings is crucial for enhancing model performance. After combining these 6 embeddings together, the t-SNE showcased clearer separation on coding RNA and lncRNA groups (Figure 3). This results in further validity of the necessary concatenate layer in LncPNdeep.

**Figure 2.**
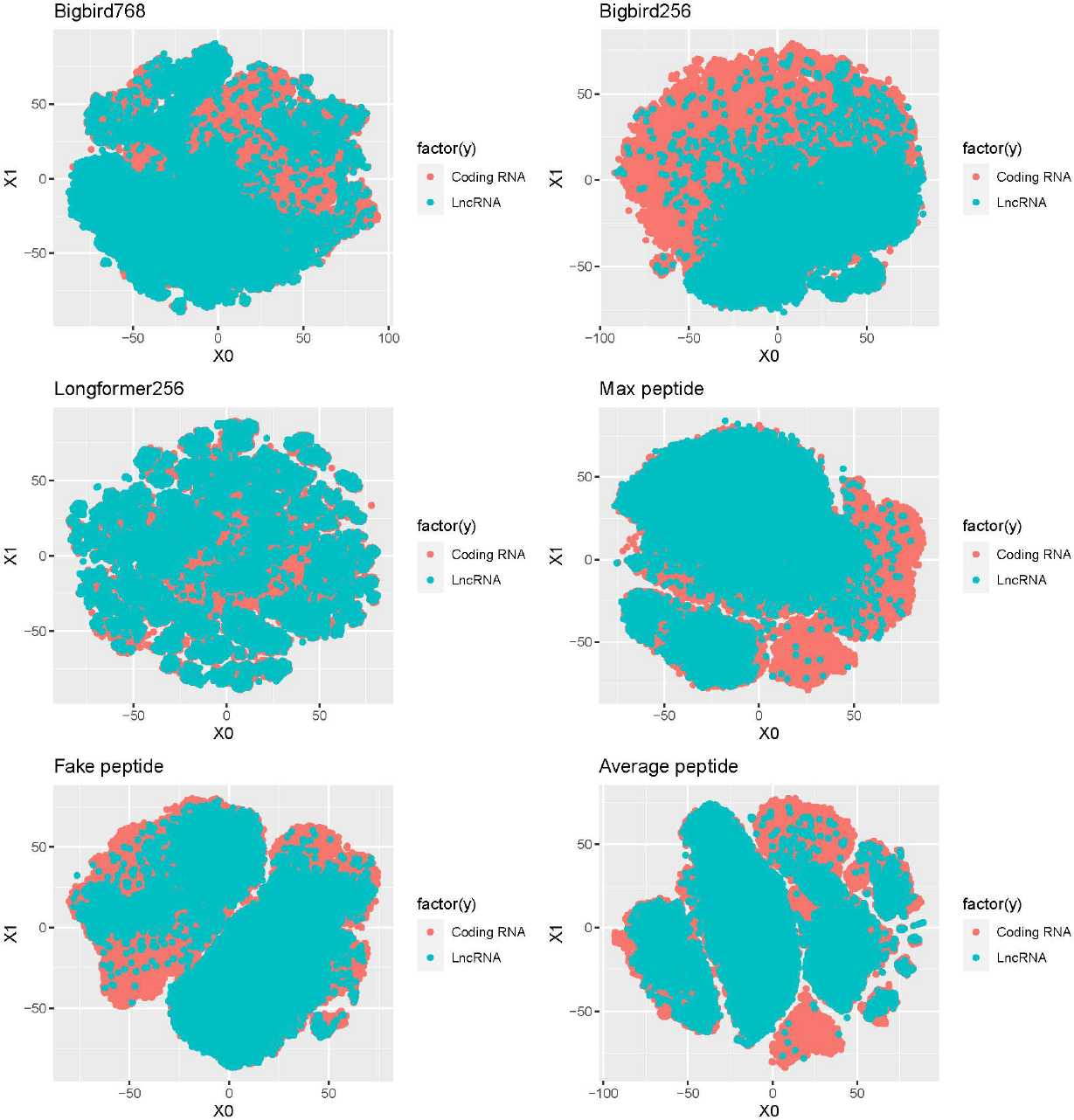
t-SNE cluster of each embedding in human RNA training dataset.

**Figure 3.**
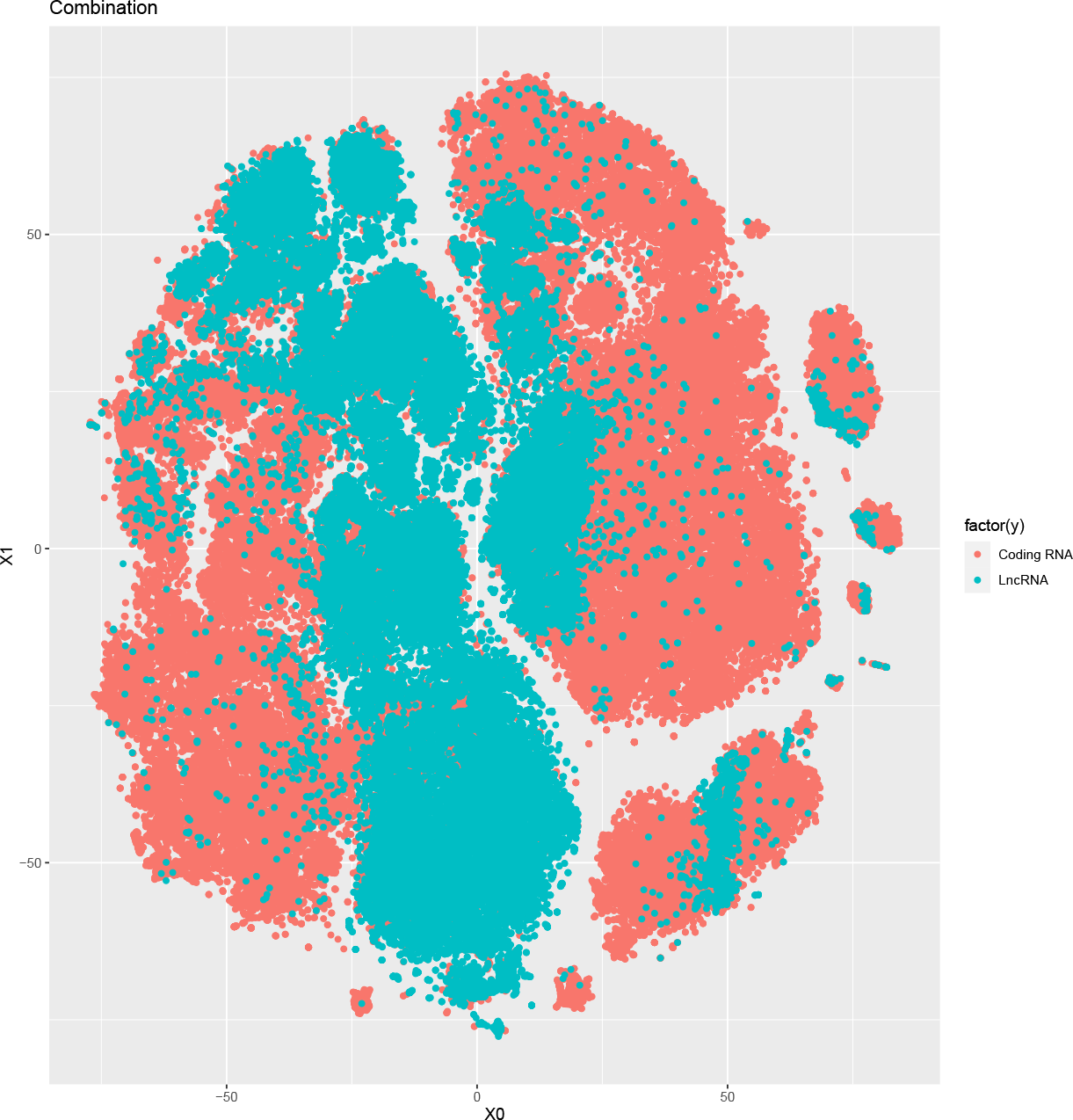
t-SNE cluster of the combinational embeddings in human RNA training dataset.

## 4. Discussion

LncRNA classification is vital for exploring the function and regulation of transcripts. Current methods relied heavily on the nucleotide features only and failed to consider the potential peptide information from RNAs. To address this problem, we designed LncPN-deep model to integrate the nucleotide and peptide information based on Concatenated Masked Language Models. This approach allows LncPNdeep to distill complex nucleotide and peptide information for more nuanced classification, providing a groundbreaking perspective in lncRNA research.

Our results indicated that LncPNdeep was a state-of-the-art model in lncRNA identification. It achieved the best performance on the human RNA database compared to the baseline model and existing methods. Furthermore, LncPNdeep also showcased remarkable generalization ability and robust performance on the cross-species dataset.

Although LncPNdeep showed superior performance on each dataset, there were some issues that needed further consideration. First, LncPNdeep utilized the Bigbird, Longformer, and ProteinTrans models for the embedding’s extraction. However, other Masked Language Models such as ProteinBERT and DNABERT remain to be assessed for potential improvement in LncPNdeep [40,41]. An optimal selection process for these models was possible to further enhance LncPNdeep’s efficacy. Second, the incorporation of protein secondary structure information, such as alpha helices and beta sheets, could yield additional insights into lncRNA classification. For poorly structured peptides, the hypothetical chemical bonds may become ill-formed and unstable, indicating special information for poor coding ability.

## Author Contributions

Conceptualization and experiment design, Zongrui Dai; Software and methodology: Zongrui Dai and Feiyang Deng; Formal analysis and data curation: Zongrui Dai and Feiyang Deng; writing—original draft preparation: Zongrui Dai and Feiyang Deng; writing—review and editing: Zongrui Dai and Feiyang Deng; visualization: Zongrui Dai

## Abbreviations

The following abbreviations are used in this manuscript:

LncRNAs: Long non-coding RNA
SVM: Support Vector Machine
CPAT: Coding-Potential Assessment Tool
MLCDS: Linear Most-Like Coding Domain Sequence
LSTM: Long short-term memory network
CNN: Convolution neural network
DSN: Deep stakced neural network
sORFs: Short open reading frames

## Disclaimer/Publisher’s Note

The statements, opinions and data contained in all publications are solely those of the individual author(s) and contributor(s) and not of MDPI and/or the editor(s). MDPI and/or the editor(s) disclaim responsibility for any injury to people or property resulting from any ideas, methods, instructions or products referred to in the content.

## References

1. Fatica, A.; Bozzoni, I. Long non-coding RNAs: new players in cell differentiation and development. Nature Reviews. Genetics 2014, 15, 1.

2. Lee, J.T.; Bartolomei, M.S. X-inactivation, imprinting, and long noncoding RNAs in health and disease. Cell 2013, 152, 1308–1323.

3. Penny, G.D.; Kay, G.F.; Sheardown, S.A.; Rastan, S.; Brockdorff, N. Requirement for Xist in X chromosome inactivation. Nature 1996, 379, 131–137.

4. Mancini-Dinardo, D.; Steele, S.J.; Levorse, J.M.; Ingram, R.S.; Tilghman, S.M. Elongation of the Kcnq1ot1 transcript is required for genomic imprinting of neighboring genes. Genes Dev. 2006, 20, 1268–1282.

5. Wang, K.C. et al. A long noncoding RNA maintains active chromatin to coordinate homeotic gene expression. Nature 2011, 472, 120–124.

6. Gupta, R.A. et al. Long non-coding RNA HOTAIR reprograms chromatin state to promote cancer metastasis. Nature 2010, 464, 7291.

7. Jia, M. et al. lincRNA-p21 inhibits invasion and metastasis of hepatocellular carcinoma through Notch signaling-induced epithelial-mesenchymal transition. Hepatology Research 2016, 46, 11.

8. Lukiw, W.J. et al. BC200 RNA in normal human neocortex, non-Alzheimer dementia (NAD), and senile dementia of the Alzheimer type (AD). Neurochemical Research 1992, 17, 6.

9. Kang, Y.J. et al. CPC2: a fast and accurate coding potential calculator based on sequence intrinsic features. Nucleic Acids Research 2017, 45, W1.

10. Li, A.; Zhang, J.; Zhou, Z. PLEK: a tool for predicting long non-coding RNAs and messenger RNAs based on an improved k-mer scheme. BMC Bioinformatics 2014, 15, 311.

11. Wang, L. et al. CPAT: Coding-Potential Assessment Tool using an alignment-free logistic regression model. Nucleic Acids Research 2013, 41, 6.

12. Sun, L. et al. Utilizing sequence intrinsic composition to classify protein-coding and long non-coding transcripts. Nucleic Acids Research 2013, 41, 17.

13. Wucher, V. et al. FEELnc: a tool for long non-coding RNA annotation and its application to the dog transcriptome. Nucleic Acids Research 2017, 45, 8.

14. Dai, Z. A New method of LncRNA classification based on ensemble learning. Journal of Physics: Conference Series 2021, 1994, No. 1, p. 012002.

15. Camargo, A.P.; Sourkov, V.; Pereira, G.A.G.; Carazzolle, M.F. RNAsamba: neural network-based assessment of the protein-coding potential of RNA sequences. NAR Genomics and Bioinformatics 2020, 2, qz024.

16. Gupta, S.; Sharma, N.K.; Shankar, R. DeepPlnc: Bi-modal deep learning for highly accurate plant lncRNA discovery. Genomics 2022, 114, 5.

17. Meng, J.; Kang, Q.; Chang, Z. et al. PlncRNA-HDeep: plant long noncoding RNA prediction using hybrid deep learning based on two encoding styles. BMC Bioinformatics 2021, 22 (Suppl 3), 242.

18. Fan, X.N.; Zhang, S.W.; Zhang, S.Y.; Ni, J.J. lncRNA_Mdeep: An Alignment-Free Predictor for Distinguishing Long Non-Coding RNAs from Protein-Coding Transcripts by Multimodal Deep Learning. International Journal of Molecular Sciences 2020.

19. Fan, X.N.; Zhang, S.W. lncRNA-MFDL: identification of human long non-coding RNAs by fusing multiple features and using deep learning. Molecular bioSystems 2015, 11, 3.

20. Ji, Z. et al. Many lncRNAs, 5’UTRs, and pseudogenes are translated and some are likely to express functional proteins. eLife 2015, 4, e08890.

21. Galindo, M.I.; Pueyo, J.I.; Fouix, S.; Bishop, S.A.; Couso, J.P. Peptides encoded by short ORFs control development and define a new eukaryotic gene family. PLoS Biol. 2007.

22. Kondo, T.; Plaza, S.; Zanet, J.; Benrabah, E.; Valenti, P.; Hashimoto, Y.; Kobayashi, S.; Payre, F.; Kageyama, Y. Small peptides switch the transcriptional activity of Shavenbaby during Drosophila embryogenesis. Science 2010.

23. Magny, E.G.; Pueyo, J.I.; Pearl, F.M.; Cespedes, M.A.; Niven, J.E.; Bishop, S.A.; Couso, J.P. Conserved regulation of cardiac calcium uptake by peptides encoded in small open reading frames. Science 2013.

24. Hartford, C.C.R.; Lal, A. When long noncoding becomes protein coding. Molecular and Cellular Biology 2020.

25. Zhang, L.; Qin, X.; Liu, M.; Liu, G.; Ren, Y. BERT-m7G: a transformer architecture based on BERT and stacking ensemble to identify RNA N7-Methylguanosine sites from sequence information. Computational and Mathematical Methods in Medicine 2021.

26. Danilevicz, M.F.; Gill, M.; Tay Fernandez, C.G.; Petereit, J.; Upadhyaya, S.R.; Batley, J.; Bayer, P.E. DNABERT-based explainable lncRNA identification in plant genome assemblies. bioRxiv 2022.

27. Brown, T.; Mann, B.; Ryder, N.; Subbiah, M.; Kaplan, J.D.; Dhariwal, P.; Amodei, D. Language models are few-shot learners. Advances in Neural Information Processing Systems 2020, 33, 1877–1901.

28. Gao, T.; Fisch, A.; Chen, D. Making pre-trained language models better few-shot learners. arXiv preprint arXiv:2012.15723 2020.

29. Liu, P.; Yuan, W.; Fu, J.; Jiang, Z.; Hayashi, H.; Neubig, G. Pre-train, prompt, and predict: A systematic survey of prompting methods in natural language processing. ACM Computing Surveys 2023, 55(9), 1–35.

30. Wang, S.; Khabsa, M.; Ma, H. To pretrain or not to pretrain: Examining the benefits of pretraining on resource rich tasks. arXiv preprint arXiv:2006.08671 2020.

31. Devlin, J.; Chang, M.W.; Lee, K.; Toutanova, K. Bert: Pre-training of deep bidirectional transformers for language understanding. arXiv preprint arXiv:1810.04805 2018.

32. Elnaggar, A. et al. ProtTrans: Toward Understanding the Language of Life Through Self-Supervised Learning. IEEE

33. Frankish, A. et al. GENCODE reference annotation for the human and mouse genomes. Nucleic Acids Research 2019, 47, D1, D766–D773. doi:10.1093/nar/gky955.

34. Martin, F.J. et al. Ensembl 2023. Nucleic Acids Research 2023, 51, D1, D933–D941. doi:10.1093/nar/gkac958.

35. Zaheer, M.; Guruganesh, G.; Dubey, K.A.; Ainslie, J.; Alberti, C.; Ontanon, S.; Ahmed, A. Big bird: Transformers for longer sequences. Advances in Neural Information Processing Systems 2020, 33, 17283–17297.

36. Beltagy, I.; Peters, M.E.; Cohan, A. Longformer: The long-document transformer. arXiv preprint arXiv:2004.05150 2020.

37. Kitaev, N.; Kaiser, Ł.; Levskaya, A. Reformer: The efficient transformer. arXiv preprint arXiv:2001.04451 2020.

38. Ainslie, J.; Ontanon, S.; Alberti, C.; Cvicek, V.; Fisher, Z.; Pham, P.; Yang, L. ETC: Encoding long and structured inputs in transformers. arXiv preprint arXiv:2004.08483 2020.

39. Child, R.; Gray, S.; Radford, A.; Sutskever, I. Generating long sequences with sparse transformers. arXiv preprint arXiv:1904.10509 2019.

40. Brandes, N.; Ofer, D.; Peleg, Y.; Rappoport, N.; Linial, M. ProteinBERT: a universal deep-learning model of protein sequence and function. Bioinformatics 2022, 38, Issue 8, Pages 2102–2110.

41. Ji, Y.; Zhou, Z.; Liu, H.; Davuluri, R.V. DNABERT: pre-trained Bidirectional Encoder Representations from Transformers model for DNA-language in genome. Bioinformatics 2021, 37, Issue 15, Pages 2112–2120.

